# Cyclin F-Chk1 synthetic lethality mediated by E2F1 degradation

**DOI:** 10.1101/509810

**Authors:** Kamila Burdova, Hongbin Yang, Roberta Faedda, Samuel Hume, Daniel Ebner, Benedikt M Kessler, Iolanda Vendrell, David H Drewry, Carrow I Wells, Stephanie B Hatch, Vincenzo D’Angiolella

## Abstract

Cyclins are central engines of cell cycle progression when partnered with Cyclin Dependent Kinases (CDKs). Among the different cyclins controlling cell cycle progression, cyclin F does not partner with a CDK, but forms an E3 ubiquitin ligase, assembling through the F-box domain, an Skp1-Cul1-F-box (SCF) module. Although multiple substrates of cyclin F have been identified the vulnerabilities of cells lacking cyclin F are not known. Thus, we assessed viability of cells lacking cyclin F upon challenging cells with more than 200 kinase inhibitors. The screen revealed a striking synthetic lethality between Chk1 inhibition and cyclin F loss. Chk1 inhibition in cells lacking cyclin F leads to DNA replication catastrophe. The DNA replication catastrophe depends on the accumulation of E2F1 in cyclin F depleted cells. We observe that SCF^cyclin F^ promotes E2F1 degradation after Chk1 inhibitors in a CDK dependent manner. Thus, Cyclin F restricts E2F1 activity during cell cycle and upon checkpoint inhibition to prevent DNA replication stress. Our findings pave the way for patient selection in the clinical use of checkpoint inhibitors.

## Introduction

Cell cycle transitions are operated by the periodic oscillations of cyclins, which bind Cyclin Dependent Kinases (CDKs) to promote phosphorylation of target substrates and drive cell cycle progression. Entry into the G1 phase of cell cycle is initiated by the activation of the transcription factor E2F1 that promotes accumulation of S phase cyclins and induces the transcription of genes required for DNA replication. The activity of E2F1 is restrained principally by the Retinoblastoma (Rb) protein, which masks E2F1 transactivation domain to keep it inactive. The control of Rb is crucial to prevent unscheduled cell cycle entry, a hallmark of cancer. Indeed, the loss of Rb is a common cancer event (Dyson, 2016).

Among the cyclins co-ordinating cell cycle progression, cyclin E/A and cyclin B in partnership with CDK2 and CDK1, respectively, promote the progression of cells through S phase and G2 phase (Lim & Kaldis, 2013). Cyclin F is most similar to cyclin A but does not act as an activator of CDKs and is not able to bind to CDKs (Bai et al, 1994). Instead, cyclin F, also known as F-box only protein 1 (Fbxo1), is the founding member of the F-box family of proteins (Bai et al, 1996). Cyclin F, through the F-box domain, forms a functional Skp1-Cul1-F-Box (SCF) complex acting as an E3 ubiquitin ligase. The SCF^cyclin F^ mediates the ubiquitylation and degradation of proteins important for cell cycle progression and genome stability (D’Angiolella et al, 2013).

In addition to the coordinated action of cyclins, cell cycle progression is monitored by the presence of checkpoints, which restrict CDK activity and prevent the execution of cell cycle phases if the previous one has not been completed.

Blocks in the progression of DNA replication fork promote the accumulation of single strand DNA (ssDNA) behind the fork, which gets coated with the ssDNA binding protein RPA, a stimuli for the activation of the ATR kinase. Active ATR elicits a checkpoint response by phosphorylating Chk1 and initiating the signalling cascade that culminates with CDK inactivation. Chk1 exerts this function by controlling the ubiquitylation and subsequent degradation of Cdc25A phosphatase (Busino et al, 2003; Jin et al, 2003; Kotsantis et al, 2018). Cdc25A removes the inhibitory phosphorylation on Tyr15 of CDKs, mediated by the tyrosine kinase Wee1 to prompt CDK activation. Upon activation of Chk1 the degradation of Cdc25A is initiated and CDK activity is suppressed to restrain DNA replication and progression of the cell cycle (Bartek & Lukas, 2003). The checkpoint response allows DNA repair before committing to mitosis. If the fork blocks cannot be bypassed or repaired, the ssDNA gaps can be converted into Double Strand Breaks (DSBs), which are more deleterious and could lead to gross chromosomal rearrangements resulting in cell death. Given their high proliferative capacity and compromised DNA repair, cancer cells have intrinsically higher replication stress and DNA damage. These observations have spurred the development of a number of inhibitors targeting the checkpoint response mediated by ATR and Chk1. Although ATR and Chk1 inhibitors are being evaluated in multiple clinical trials, the genetic predispositions that would sensitise or desensitise cancer cells to ATR and Chk1 inhibitors are not fully elucidated.

By screening the Kinase ChemoGenomic Set (KCGS) to identify vulnerabilities of cells lacking cyclin F, we discovered a novel synthetic lethal interaction between cyclin F loss and Chk1 inhibition. We observed that the loss of cyclin F promotes DNA replication catastrophe after challenging cells with Chk1 inhibitors. The DNA replication catastrophe in cells lacking cyclin F after Chk1 inhibitors depends on the accumulation of E2F1. E2F1 is a novel substrate of cyclin F which is controlled at the G2/M transition and after checkpoint inhibition. Analogous to the DNA replication stress induced by oncogene activation (Kotsantis et al, 2018), E2F1 accumulation in cyclin F knockout cells exacerbates Chk1 inhibitor sensitivity, promoting the accumulation of DSBs and cell death.

## Results

### KCGS screen identifies synthetic lethality between Chk1 inhibitors and cyclin F KnockOut (K/O)

Using CRISPR (Clustered Regularly Interspaced Short Palindromic Repeats)/Cas9 we have previously generated cyclin F KnockOut (*CCNF* K/O) cell lines (Mavrommati et al, 2018). Interestingly, these cells present a cell cycle profile comparable to the control parental cell line (Figure EV1A and B). To identify weaknesses of cells lacking cyclin F, we conducted a kinase inhibitor drug screen to assess differences of viability between *CCNF* K/O and parental cell lines (HeLa) (Figure 1A). To this end, we used the KCGS, a set whose origins can be traced to the well-utilized kinase inhibitor collections PKIS and PKIS2 (Drewry et al, 2017; Elkins et al, 2016). The simple premise of KCGS is that screening a publicly available, well annotated set of kinase inhibitors in disease relevant phenotypic assays is an efficient way to elucidate biology and uncover dependencies (Jones & Bunnage, 2017). The current version of KCGS consists of 188 small molecule kinase inhibitors that cover 221 kinases. None of the members of KCGS is promiscuous, facilitating target deconvolution. The screen was run in duplicate with an average replicate correlation (Pearsons’) of 0.9467 and an average Z-Factor of 0.6040, demonstrating a good dynamic range between DMSO (vehicle negative control) and controls (chemo-toxic positive control) with robust reproducibility between replicates. To identify small molecule kinase inhibitors, which specifically target cyclin F depleted cells, we normalized the cell viability results using a Z Score for each cell line and then expressed the difference between *CCNF* K/O cells and control cells as ΔZ. The graph in Figure 1A and additional data in Table 1 highlight compounds to which *CCNF* K/O cells are sensitive or resistant. The most striking difference in viability between control cells and *CCNF* K/O cells was obtained with CCT244747, a selective Chk1 kinase inhibitor (Walton et al, 2012). Among the compounds, which were killing *CCNF* K/O cells, we also identified VE-822, an ATR inhibitor (Charrier et al, 2011). Similarly, in a parallel drug screen a striking sensitivity of *CCNF* K/O to two other Chk1 inhibitors with unrelated structure, PF477736 and AZD7762, was observed (Figure 1B)(Blasina et al, 2008; Zabludoff et al, 2008).

**Figure 1.**
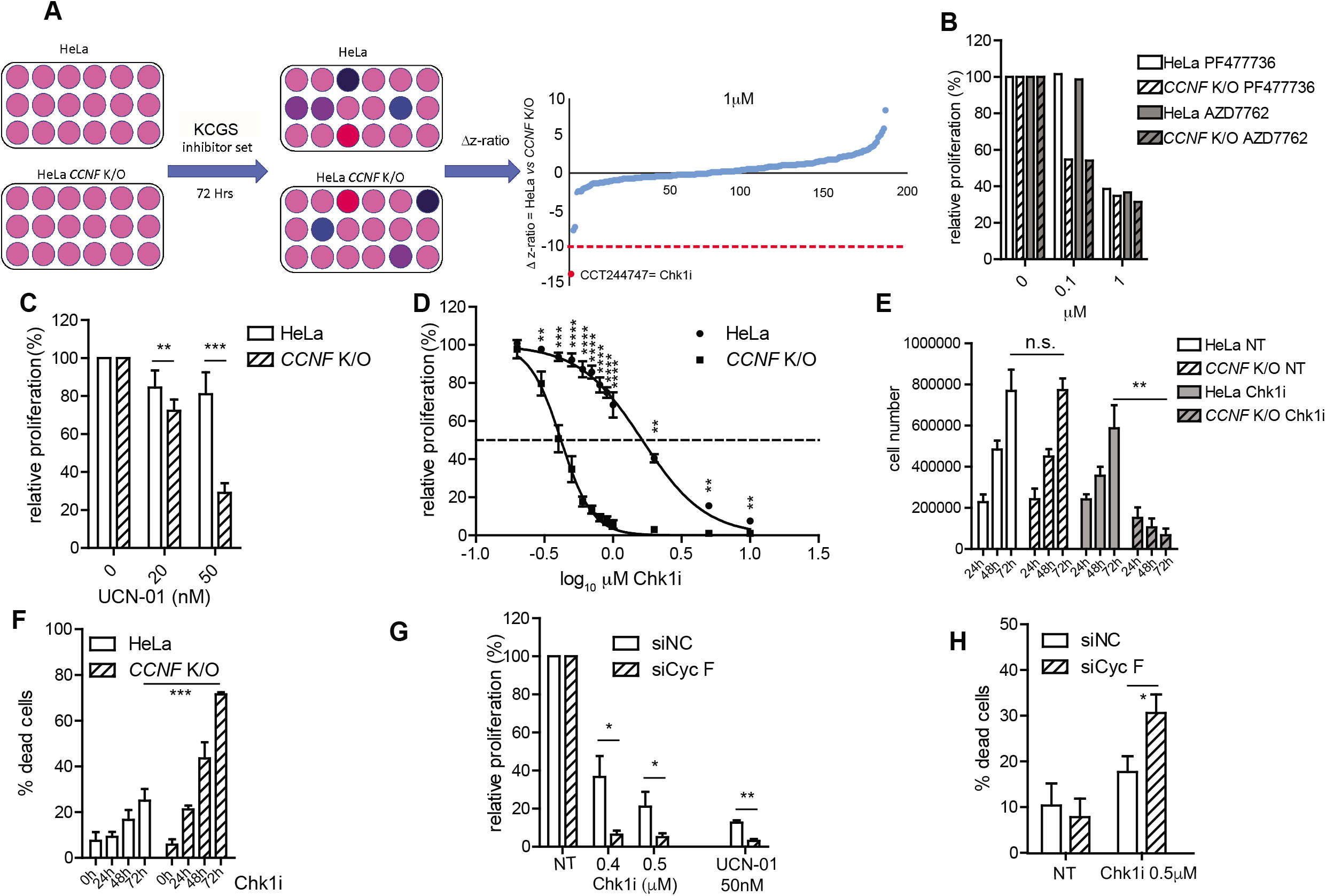
KCGS screen identifies synthetic lethality between Chk1 inhibitors and cyclin F KnockOut (K/O) A. HeLa and cyclin F KnockOut (*CCNF* K/O) cells were treated with Kinase ChemoGenomic Set (KSGS) in 384 well format. After 72 Hours (h) viability was measured using rezasurin. Flowchart representation (*left*). Robust Z-score difference is plotted for data acquired at 1 μM concentration (*right*). B. Cell survival measured using rezasurin and compared to controls treated with DMSO (expressed as relative proliferation %). Cells were treated with the indicated inhibitors at 0.1 and 1 μM concentration. C. Cell survival measured using rezasurin and compared to controls (0) treated with DMSO (expressed as relative proliferation %). Cells were treated with UCN-01 at the indicated concentrations. For all bar graphs presented henceforth, data are presented as mean ± standard deviation (SD), with at least 3 independent experiments. p-values (* p<0.05, ** p<0.005, *** p<0.0005, ****p<0.00005) were calculated by paired and two-tailed t test. NT= treatment with DMSO. D. Differences in viability of HeLa and *CCNF* K/O cells after treatment with specific Chk1i (LY2603618) at the indicated concentrations plotted on a log_10_ scale to measure differences in IC50. E. Number of cells was measured in HeLa and *CCNF* K/O cells left untreated or treated with Chki (LY2603618) for 24h, 48h, or 72h as indicated. (n.s.= non significant). F. Cell viability measurements using Propidium Iodide staining for HeLa and *CCNF* K/O cells after treatment with Chk1i (LY2603618). G. U-2-OS cells were seeded and transfected with non targeting siRNA siNC (Negative Control) or siCyc F. 24 hours after transfection, cells were challenged with UCN-01, or Chk1i (LY2603618) at indicated concentrations for 3 days before viability was measured. H. Cell viability measurements using Propidium Iodide staining in U-2-OS cells transfected with non-targeting siNC and siCyc F and treated with Chk1i (LY2603618) as indicated.

**Table 1.** Top hits of the viability screen comparing HeLa to *CCNF* K/O using the KCGS.

| 100nM | | compound name | $\Delta$ z-score | Target |
| --- | --- | --- | --- | --- |
| more sensitive | C10 | VE-822 | -0.61484 | ATR |
|  | B17 | CCT251545 | -0.56548 | CDK8, CDK19 |
|  | I03 | GSK2336394 | -0.4252 | PIK3CB |
| less sensitive | A20 | GW784752X | 1.016439 | GSK3b |
|  | A21 | GW580509X | 1.052113 | VEGFR2 |
|  | E22 | GW296115X | 1.443446 | PDGFR |
| 1μM | | compound name | $\Delta$ z-score | Target |
| more sensitive | B08 | CCT244747 | -2.88143 | CHK1 |
|  | E16 | BI00614644 | -0.56505 | MAPKAPK2, MAPKAPK5, PIM3, PHKG2 |
|  | I05 | SB-725317 | -0.55738 | CDK2/cyclinA, LTK |
| less sensitive | C11 | SB-590885 | 1.036447 | B-raf |
|  | B03 | JNK-IN-7 | 1.164645 | JNK |
|  | E22 | GW296115X | 1.180556 | PDGFRB |
|  | D15 | GW814408X | 1.364238 | GSK3b |
|  | H16 | GSK1070916 | 1.572064 | Aurora kinase |
|  | D12 | CA93.0 | 2.072805 | GAK1 |
| 10μM | | compound name | $\Delta$ z-score | Target |
| more sensitive | H19 | GW807982X | -1.06323 | GSK3b |
|  | I20 | TPKI-39 | -0.86181 | CIT, KIT, PDGFRB, CSF1R, PDGFRA, DDR1, FLT1, TIE2 |
|  | C13 | GSK1838705 | -0.73301 | IGF1R, IR |
| less sensitive | G14 | PFE-PKIS 3 | 0.838916 | TNK2, TYRO3, YES, FGFR1,2,&3 |
|  | H11 | CCT241533 | 0.859239 | CHK2 |
|  | G09 | GW440135 | 0.866643 | KIT, NEK7, MEK5, PDGFRB, VEGFR2, NEK9, PDGFRA, FLT4 |

To confirm that the effect of CCT244747, PF47736 and AZD7762 was on target we tested survival of HeLa *CCNF* K/O also with LY2603618 and UCN-01, two unrelated widely used Chk1 inhibitors. UCN-01 decreased cell proliferation in *CCNF* K/O cells compared to HeLa parental cells (Figure 1C). Upon treatment of cells with LY2603618, a very selective Chk1 inhibitor (King et al, 2014), we observed the most striking difference in cell viability. The IC50 of LY2603618 in normal HeLa cells was > 1 uM, whereas *CCNF* K/O cells had an IC50 of 400 nM, accounting for more than several fold difference in sensitivity across the two cell lines using LY2603618 (Figure 1D).

To establish that the difference in sensitivity between control cells and *CCNF*K/O was not due to proliferation defects of *CCNF* K/O cells, we measured cell proliferation across 72 hours comparing HeLa control to *CCNF* K/O. Overall cell proliferation was not different in the 72 hours time-frame between HeLa control and *CCNF* K/O (Figure 1E). However, upon addition of LY2603618, *CCNF* K/O cells stopped proliferating (Figure 1E), corresponding to an increase in cell death (Figure 1F).

To exclude the possibility that the phenotype of cell death induced by Chk1 inhibitors was limited to *CCNF* K/O cell lines, we also tested sensitivity of U-2-OS cells to LY2603618 after knockdown of cyclin F by siRNA. Although treatment of U-2-OS cells with LY2603618 at 500 nM caused more cell death in U-2-OS cells compared to HeLa cells, U-2-OS cells where cyclin F levels were diminished by siRNA, had significantly reduced cell survival and increased cell death (Figure 1G & H).

Since ATR is upstream of Chk1, it was not surprising that the drug screen above also identified ATR inhibitors. Indeed, inhibition of ATR using VE-821, also promoted cell killing in *CCNF* K/O, although the difference between HeLa control and *CCNF* K/O was only observed at high doses and was less pronounced compared to Chk1 inhibitors (Figure EV1C). Similar results were obtained after inhibition of ATR in U-2-OS cells and cyclin F depletion by siRNA (Figure EV1D). The data presented above highlight a novel synthetic lethal interaction between the loss of cyclin F and checkpoint depletion. The sensitivity of cyclin F loss to Chk1 inhibition was striking and investigated further.

### Loss of cyclin F causes DNA replication catastrophe after Chk1 inhibition

The inhibition of ATR and Chk1 promotes accumulation of ssDNA, which is exacerbated over time due to active DNA replication. The ssDNA saturates the capacity of RPA to bind to it and is exposed to the action of nucleases. Nucleases convert ssDNA to DSBs, which promote the activation of DNA-PK and ATM kinases required for DSB repair. The accumulation of DSBs can be detected by markers of DNA-PK and ATM kinase activation and is an indication of DNA replication catastrophe (Toledo et al, 2013).

To gain insights into the mechanism of synthetic lethality between loss of cyclin F and Chk1 inhibition, we compared the response of HeLa parental cells and *CCNF* K/O cells after treatment with Chk1 inhibitor. In contrast to parental cells that showed accumulation of RPA phosphorylation on Serine (S) 33, a surrogate marker of ATR activation, at 6 hours, *CCNF* K/O cells already showed increased RPA S33 phosphorylation after 2 hours of treatment. While ATR activation can be attributed to accumulation of RPA-coated ssDNA, further activation of DNA-PK and ATM can be associated with formation of DSBs leading to DNA replication catastrophe detected by phosphorylation of γ-H2AX, phosphorylation of RPA on S4 and S8 and phosphorylation of KAP1 on S824 at the 16-hour time point as reported previously (Toledo et al, 2013). In *CCNF* K/O cell line, phosphorylation of γ-H2AX, phosphorylation of RPA on S4 and S8 and phosphorylation of KAP1 on S824 were already present at 6 hours after treatment with Chk1 inhibitor (LY2603618). Activation of the DSB markers continued thereafter, accumulating profoundly at 16 and 24 hours of treatment compared to control cells (Figure 2A). It is worth mentioning that the overall levels of Chk1 and RPA were unchanged in *CCNF* K/O cells compared to control cells, indicating that the differences observed were due to upstream phosphorylation events rather than a change in the total levels of these proteins (Figure 2A). Similar results were obtained in U-2-OS cell line using siRNA knock-down of cyclin F after treatment with Chk1 inhibitor (Figure EV2A).

**Figure 2.**
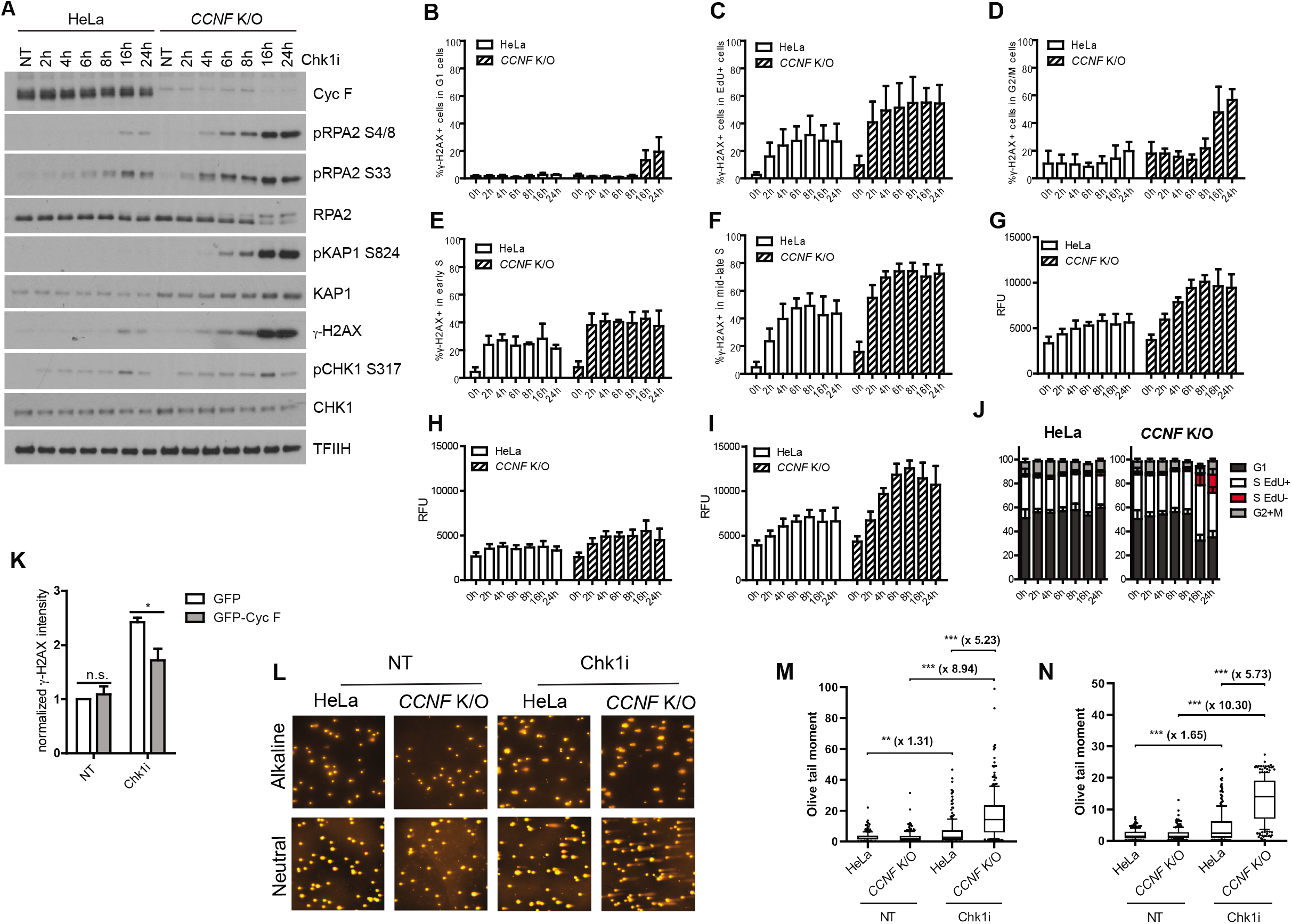
Loss of cyclin F causes DNA replication catastrophe after Chk1 inhibition. A. Cells treated with 1μM Chk1i (LY2603618) for the indicated hours (h) were harvested and lysed using SDS. Indicated protein were resolved by SDS page and detected by Western blot (Wb). TFIIH was used as a loading control. B. Percentage (%) of γ-H2AX positive cells in G1 cells after Chk1i (LY2603618) treatment for the indicated hours (h) measured using Fluorescence Activated Cell Sorter (FACS). G1 cells were considered having DAPI 2n staining and EdU negativity. Unless otherwise stated, all FACS data shown henceforth includes data from at least 3 independent biological replicates and data were plotted with Mean ±SD. C. Percentage (%) of γ-H2AX positive cells in replicating (EdU+) cells after Chk1i (LY2603618) treatment at the indicated time points. Data were plotted with Mean %± SD. D. Percentage (%) of γ-H2AX positive cells in G2/M cells after Chk1i (LY2603618) treatment. G2/M cells were considered having DAPI 4n staining and EdU negativity. Data were plotted with Mean % ± SD. E. Percentage (%) of γ-H2AX positive cells in early S cells after Chk1i treatment (LY2603618). Early S phase cells were considered having 2n DAPI content and but EdU+. Data were plotted with Mean %± SD. F. Percentage (%) of γ-H2AX positive cells in mid-late S phase after Chk1i (LY2603618) treatment. Mid-late S phase cells were considered having DAPI 4n staining and EdU+. Data were plotted with Mean % ± SD. G. Intensity of γ-H2AX measured as Relative Fluorescence Units (RFU) in replicating cells treated with Chk1i. Mean intensity of γ-H2AX fluorescence was evaluated from EdU+. Data were plotted with Mean RFU ± SD. H. Intensity of γ-H2AX measured as Relative Fluorescence Units (RFU) in early S phase cells treated with Chk1i. Mean intensity of γ-H2AX fluorescence was evaluated from EdU+ cells with 2N DAPI content. Data were plotted with Mean RFU ± SD. I. Intensity of γ-H2AX measured as Relative Fluorescence Units (RFU) in mid-late S phase cells treated with Chk1i. Mean intensity of γ-H2AX fluorescence was evaluated from EdU+ cells with > 2N DAPI content. Data were plotted with Mean RFU ± SD. J. Overall, cell cycle profile of HeLa and *CCNF* K/O cells after Chk1i treatment. Cells with 2n DAPI staining but EdU-were gated as G1 cells; cells with >2n but <4n DAPI staining but EdU-were gated as EdU-S cells; cells with >2n but <4n DAPI staining but EdU+ were gated as EdU+ S phase; cells with 4n DAPI and pH3 S10 negative staining were gated as G2 phase and cells with 4n DAPI and pH3 S10 positive as M cells. K. Normalised γ-H2AX intensity in *CCNF* K/O cells transfected with either GFP empty vector or GFP-Cyc F and treated with 1μM Chk1i for 20 hours. GFP positive cells were gated. Data were plotted with Mean % ± SD. Related to Supplementary Figure 2C. L. Representative images of neutral and alkaline comet assay from HeLa and *CCNF* K/O cells treated with Chk1i. Cells were treated with 1μM Chk1i for 24h before being harvested for neutral and alkaline comet assays as indicated. M. Quantification of comets in <u>***alkaline***</u> gels. At least 100 cells, across two slides, were analysed in each condition in two biological replicates. Data are shown as medians, with 25/75% percentile range (box) and 10-90% percentile range (whiskers). Fold changes of median values are shown in bold. p-values were calculated using the Mann-Whitney test (two-tailed). Olive tail moment = (Tail.mean - Head.mean)*Tail%DNA/100. N. Quantification of comets in <u>***neutral***</u> gels. At least 100 cells, across two slides, were analysed in each condition in two biological replicates. Data are shown as medians, with 25/75% percentile range (box) and 10-90% percentile range (whiskers). Fold changes of median values are shown in bold. p-values were calculated using the Mann-Whitney test (two-tailed). Olive tail moment = (Tail.mean - Head.mean)*Tail%DNA/100.

To confirm that *CCNF* K/O cells indeed encountered DNA replication catastrophe, we analysed chromatin bound RPA and γ-H2AX formation in response to Chk1 inhibitor in different cell cycle phases by Fluoresce-activated cell sorting (FACS). As expected, the γ-H2AX signal was coming from replicating cells (Figure 2B, C, D, G) in early time points and, γ-H2AX levels increased as cells progressed from early S to mid-late S phase (Figure 2E, F, H, I). Only at later time points (16-24h) did cells enter the G2 phase of the cell cycle, with an increase in γ-H2AX levels (Figure 2 B, D). The increased γ-H2AX signal was accompanied by a strong increase in RPA binding to chromatin and cell cycle arrest in S-phase (Figure EV2B). At 24h after Chk1 inhibitor treatment, about 15% of cells were found to be highly RPA and γ-H2AX positive but EdU negative, having S phase DNA content of 2-4n (Figure 2J). Most importantly, γ-H2AX formation in *CCNF* K/O cells upon Chk1 inhibitor treatment was significantly reduced after expression of GFP-cyclin F (Figure 2K and Figure EV2C), demonstrating that the formation of DSBs is mediated specifically by cyclin F loss.

Although the evidence provided so far is indicative of DNA replication catastrophe in *CCNF* K/O cells upon Chk1 inhibition, we sought direct evidence of single strand breaks and DSB accumulation. Therefore, we conducted alkaline and neutral comet assays in *CCNF* K/O cells treated with Chk1 inhibitors. Treatment of *CCNF* K/O cells with Chk1 inhibitors induced a drastic increase in single strand breaks and DSBs readily visible in the tail images (Figure 2L) and quantified in the olive tail moment (Figure 2M and N). Our data support a model where loss of cyclin F predisposes cells to DNA replication catastrophe upon challenging cells with Chk1 inhibitors.

### E2F1 mediates DNA replication catastrophe in *CCNF* K/O cells treated with Chk1 inhibitors

To identify novel substrates of cyclin F that mediate DNA replication stress after checkpoint inhibition, we performed immunoprecipitation of cyclin F and identified interacting partners by Liquid Chromatography/ Mass Spectrometry (LC/MS). A statistically significant enrichment for G1/S phase transition regulators and DNA replication processes was identified (Figure 3A; p -value <0.002). We speculated that these interacting partners likely accumulate in *CCNF* K/O cell lines and exacerbate DNA replication stress upon Chk1 inhibition. Since components of pre-RC (Cdc6, Cdt1 and Geminin), components of the DNA replication elongation machinery (TICRR and MTBP), G1/S cell cycle transcriptional regulators (E2F1 and E2F3) and exonucleases (Exo1) have all been previously linked to the induction of DNA replication stress, we focused our study on these factors (Hills & Diffley, 2014; Kotsantis et al, 2018). To seek the interacting partner of cyclin F which could mediate sensitivity to Chk1 inhibitors, we performed siRNA of Exo1, Cdc6, Cdt1, Geminin, MTBP, TICRR, E2F1 and E3F3 in HeLa control and *CCNF* K/O (Figure EV3A, B, C and D) after treatment with Chk1 inhibitors and determined the extent of DNA replication stress by measuring γ-H2AX levels and RPA accumulation on chromatin respectively by FACS. We obtained a complete rescue of γ-H2AX and RPA accumulation only after siRNA of E2F1 but not after siRNA of any factor indicated above, suggesting that E2F1 was the main determinant of Chk1 inhibitor sensitivity upon cyclin F depletion (Figure 3 B and C). A different siRNA oligo targeting E2F1 but not siRNA targeting Exo1 rescued sensitivity of *CCNF* K/O cells to Chk1 inhibitors (Figure EV3E). Notably, high expression of exogenous E2F1 in HeLa cells was sufficient to mediate DNA replication stress after Chk1 inhibition, as visualised by increased levels of γ-H2AX intensity detected by FACS analysis (Figure 3D and Figure EV3F). All the surrogate markers of ssDNA and DSB formation (phosphorylation of RPA on S33, phosphorylation of RPA on S4 and S8, γ-H2AX and phosphorylation of KAP1 at S824) were increased in *CCNF* K/O cells upon treatment with Chk1 inhibitors as observed above. However, siRNA of E2F1 was sufficient to restore the levels of these markers to basal levels present in HeLa controls (Figure 3E). A potential downstream target of E2F1, which could mediate the observed DNA replication catastrophe is cyclin E whose overexpression has been extensively associated with DNA replication stress(Ekholm-Reed et al, 2004; Macheret & Halazonetis, 2018). However, siRNA of cyclin E in *CCNF* K/O cells only partially suppressed DNA replication catastrophe (Figure EV3G), indicating the presence of other targets of E2F1 mediating DNA replication catastrophe in addition to cyclin E.

**Figure 3.**
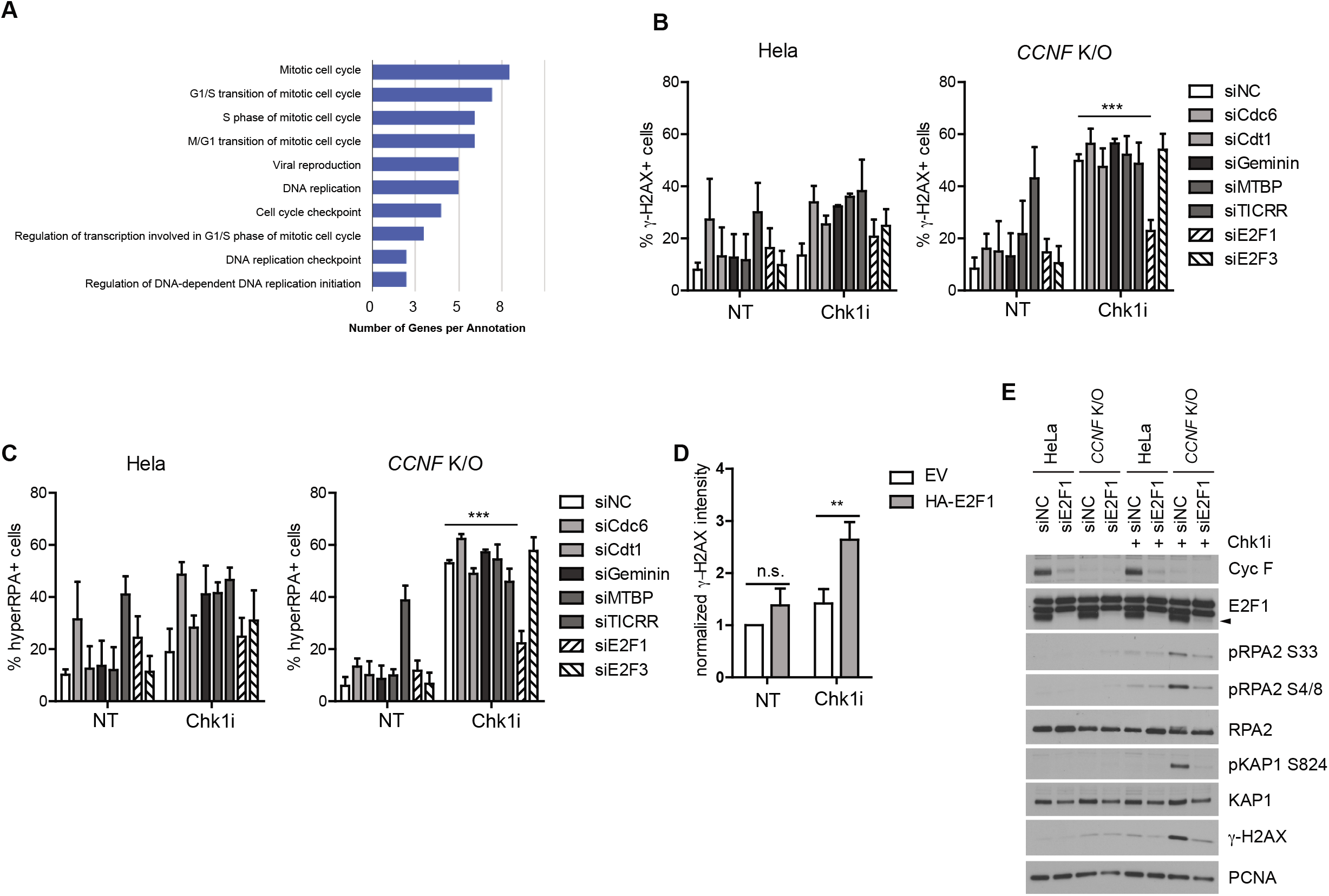
E2F1 mediates DNA replication catastrophe in cyclin F K/O cells treated with Chk1 inhibitors. A. Pathway enrichment analysis using GENECODIS of Cyclin F high confidence interacting partners identified by Liquid Chromatography/Mass Spectrometry (LC/MS) using YFP-cyclin F as bait. B. Percentage (%) of γ-H2AX positive cells in HeLa and *CCNF* K/O after transfection of the indicated siRNAs treated with Chk1i for 20 hours. Percent of γ-H2Ax positive cells were plotted as Mean % ± SD. C. Percentage (%) of hyper-RPA positive cells in HeLa and *CCNF* K/O after transfection of the indicated siRNAs treated with Chk1i for 20 hours. Percent of hyper RPA positive cells were plotted as Mean %± SD. D. Relative γ-H2AX intensity in cells transfected with either Empty Vector (EV) or HA-E2F1 treated with Chk1i for 20 hours. γ-H2AX fluorescence in HA positive cells was normalized to the empty vector transfected cells and compared to cells not expressing HA. Related to Supplementary Figure 3F. E. HeLa and *CCNF* K/O cells transfected with the indicated siRNA and treated with 1μM Chk1i (LY2603618) for 20 hours were harvested and lysed using SDS. Indicated protein were resolved by SDS page and detected by Wb. PCNA served as a loading control.

A similar set of experiments to the ones described above was conducted upon siRNA of cyclin F and treatment with ATR inhibitors. We observed DNA replication stress upon cyclin F depletion and ATR inhibition, as detected by the appearance of phosphorylation of RPA on S4, S8 and γ-H2AX. The DNA replication stress observed in cyclin F depleted cells treated with ATR inhibitor was rescued by concomitant depletion of E2F1 (Figure EV3H). Furthermore, we conducted a comet assay to detect directly DNA damage in cells treated as above. The comet assay highlighted that ATR inhibition induced increased DNA damage after siRNA of cyclin F. The DNA damage present in cells with reduced cyclin F levels was rescued fully by concomitant depletion of E2F1 (Figure EV3I). We conclude that DNA replication catastrophe induced by cyclin F depletion and checkpoint inhibition is mediated by E2F1.

### Cyclin F regulates E2F1 after Chk1 inhibition

The experiments presented above suggest that E2F1 is a novel target of cyclin F. Therefore, we asked whether E2F1 was regulated by cyclin F and measured the levels of E2F1 in *CCNF* K/O cells or after siRNA. *CCNF* K/O cell lines had significantly increased levels of E2F1 compared to control (Figure EV3B, Figure 4A). Furthermore, siRNA of cyclin F induced a significant increase of E2F1 in U-2-OS cells (Figure 4B).

**Figure 4.**
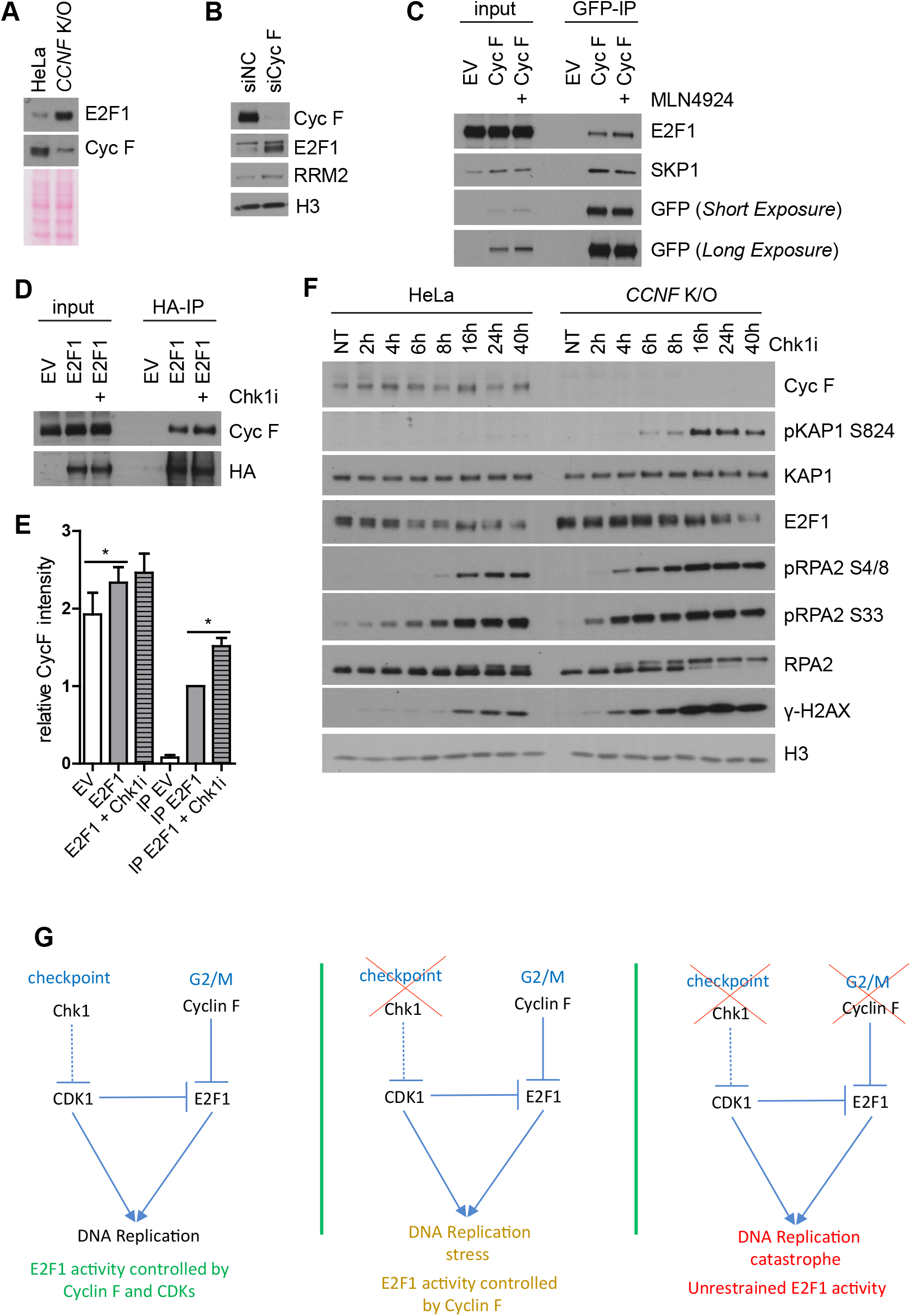
Cyclin F regulates E2F1 after Chk1 inhibition. A. HeLa and *CCNF* K/O cells were harvested and lysed using SDS. Indicated protein were resolved by SDS page and detected by Wb. Ponceau S staining was used as a loading control. B. U2OS cells transfected with non targeting siRNA (siNC) or with three siRNA targeting Cyc F were lysed using SDS. Whole cell lysates were analyzed by immunoblotting 48h after transfection. H3 was used as a loading control. C. HEK293 cells transfected with GFP empty vector (EV) or YFP-Cyclin F were treated with MLN4924 for 4 hours and harvested for immunoprecipitation using GFP-nanobodies. Immunoprecipitates were immunoblotted as indicated. D. HEK293 cells transfected with empty vector (EV) or HA-E2F1 were treated with Chk1i for 20 hours and harvested for immunoprecipitation using HA antibodies. Immunoprecipitates were immunoblotted as indicated. E. Quantification of immunoprecipitation results of Figure 4D form three independent experiments. Data were plotted with Mean % ± SD. *< 0.05 p-vaue. F. HeLa and *CCNF* K/O cells treated with 1μM Chk1i (LY2603618) for the indicated time points (h=hours) were harvested and lysed using SDS. Indicated protein were resolved by SDS page and detected by Wb. H3 served as a loading control. G. Schematic representation of a model depicting the mechanism eliciting synthetic lethality between cyclin F loss and Chk1 inhibition.

To prove that the regulation of E2F1 by cyclin F was direct, we tested by immunoprecipitation whether E2F1 could interact with cyclin F. Cyclin F could interact specifically with E2F1 and the interaction between cyclin F and E2F1 was reinforced by treating cells with MLN4924, an inhibitor of Nedd8 activating enzyme, which prevents the activity of SCF complexes (Soucy et al, 2009) (Figure 4C). The half-life of E2F1 was increased in *CCNF* K/O cells (Figure EV4A). Furthermore, the interaction between cyclin F and E2F1 was potentiated after treating cells with Chk1 inhibitors (Figure 4D and quantified in E). Finally, Chk1 inhibitors induced a reduction of E2F1 levels, which was prevented in cells lacking cyclin F (Figure 4F). We have previously shown that RRM2, another substrate of cyclin F, is ubiquitylated by cyclin F only after phosphorylation by CDKs (D’Angiolella et al, 2012). As Chk1 inhibitors promote the activation of CDKs we speculated that the phosphorylation by CDKs could be a common feature across cyclin F substrates. To this end, we tested whether phosphorylation impacted the interaction between cyclin F and E2F1. Treatment of immunoprecipitated cyclin F with a non-specific phosphatase (λ) did not change the interaction between cyclin F and E2F1 although RRM2 was released after dephosphorylation of Threonine 33, as previously reported (Figure EV4B). At the same time, treatment of cells with a serine-threonine phosphatase inhibitor (Calyculin A) strongly promoted cyclin F and E2F1 interaction (Figure EV4B). It could be speculated that E2F1 phosphorylation promotes interaction with cyclin F but phosphorylation is just a priming mechanism for interaction. To test whether the priming phosphorylation for interaction with cyclin F was mediated by CDKs, we treated cells with specific CDK1 and CDK2 inhibitors combined with calyculin A to stabilise the interaction. As expected, the interaction between cyclin F and RRM2 was abolished by treatment with CDK inhibitors. Under the same conditions, the interaction between cyclin F and E2F1 was reduced after treatment with CDK1 inhibitors and was abolished when both CDK inhibitors were present (Figure EV4C).

E2F1 levels are reduced during mitosis and G2/M when CDK activity is maximal (Budhavarapu et al, 2012). Therefore, we synchronized cells with nocodazole in prometaphase and measured the levels of E2F1 and other known substrates of cyclin F. HeLa parental cells arrested in mitosis had drastically reduced levels of E2F1 compared to asynchronous cells. On the contrary, *CCNF* K/O cells showed a significant increase in E2F1 levels and its half-life (Figure EV4D). *CCNF* K/O cells transferred to the next G1 phase of cell cycle when released from nocodazole treatment also showed increased E2F1 levels (Figure EV4E). Our data point out that E2F1 is regulated directly by cyclin F after Chk1 inhibition and during cell cycle progression.

## Discussion

Cyclin F is emerging as an important constituent of the G2/M transition that promotes the regulation of central cell cycle factors. Here, starting from a drug screen, we identify a novel synthetic lethal interaction between cyclin F loss and Chk1 inhibition. Our study highlights that *CCNF* K/O cells, upon Chk1 inhibition, undergo DNA replication catastrophe, which is followed by cell death. We observe that the synthetic lethality is mediated by the accumulation of E2F1, present in cyclin F depleted cells. The concomitant depletion of E2F1 prevents DNA replication catastrophe in *CCNF* K/O cells. We further show that E2F1 protein is regulated by cyclin F and the interaction between cyclin F and E2F1 is potentiated by Chk1 inhibitors. We propose a model where Chk1 inhibitors deregulate CDK activity to prompt DNA replication stress. In these circumstances, cyclin F can restrict E2F1 activity by controlling its levels and thus, suppressing DNA replication progression. The concomitant loss of cyclin F and Chk1 prompts uncontrolled E2F1 activity when CDK activity is unscheduled, inducing DNA replication catastrophe (Figure 4G).

The mechanism of DNA replication catastrophe induced by E2F1 accumulation in cyclin F depleted cells could be seen as oncogene induced DNA replication stress, mostly connected to models of cyclin E overexpression. It has been shown that cyclin E accumulation could lead to reduced DNA replication origin assembly (Ekholm-Reed et al, 2004). Furthermore, cyclin E overexpression alters the normal DNA replication programme forcing the formation of intragenic origins of replication. This process predisposes cells to the deleterious consequences of transcription replication collisions and results in the formation of DSBs (Macheret & Halazonetis, 2018). Our findings suggest that E2F1 mediates DNA replication stress partially through cyclin E. Thus, it is possible that the activation of E2F1 in conditions where the activity of Cyclin E/CDK2 is deregulated could further exacerbate transcription replication collisions and contribute to the induction of DNA replication catastrophe in *CCNF* K/O cells. It is important to note that the DNA replication stress induced by cyclin F depletion is different from cyclin E, as it does not rely on overexpression systems.

There is suggestive evidence that transcription and DNA replication initiation sites overlap (https://www.biorxiv.org/content/early/2018/05/16/324079), appropriate regulation of E2F1 during cell cycle could be crucial in this process. Interestingly, cyclin F regulates E2F1 only in the G2/M phase of the cell cycle. It is tantalising to speculate that cyclin F regulates a late DNA replication programme, which entails E2F1 depletion. The *CCNF* K/O system of E2F1 deregulation could resolve longstanding questions in the field of DNA replication stress and warrants further investigation.

Chk1 inhibitors are being evaluated in clinical trials in a variety of solid tumors. Inactivating mutations in cyclin F detected as diploid truncating mutations have been identified in stomach adenocarcinoma, lung adenocarcinoma, medulloblastoma, squamous cell carcinoma of the head and neck, glioblastoma, mucinous adenocarcinoma of the colon and rectum, colon carcinoma and cutaneous melanoma. In light of our observations we believe that patients should be selected based on cyclin F status. Given the striking synthetic lethality we report, patients with cancers bearing truncating mutations in cyclin F could benefit significantly from treatment with Chk1 inhibitors.

Interestingly, our drug screen reveals a number of positive and negative drug interactions, which could be further investigated. Cyclin F is amplified in 10-15% of breast invasive ductal carcinoma. It is likely that amplification of cyclin F hampers the effectiveness of Chk1 inhibitor treatment. In the future, it would be crucial to establish cancer models that mimic cyclin F amplification both *in vivo* and *in vitro*.

## Acknowledgments

This study was possible thanks to the support of a Medical Research Council (MRC) grant MC_UU_00001/7 to V. D’A. This work was supported by Cancer Research UK (CR-UK) grant number C5255/A18085, through the Cancer Research UK Oxford Centre. This work was further supported by a John Fell (133/075) and Wellcome Trust grant (097813/Z/11/Z) to B.M.K. Funding for the SGC-UNC was provided by The Eshelman Institute for Innovation, UNC Lineberger Comprehensive Cancer Center, PharmAlliance, and National Institutes of Health (1R44TR001916-02, 1R01CA218442-01, and U24DK116204-01).

## Materials and Methods

### Cell lines and synchronization

HEK293, U-2-OS, MCF7, Hela, and HeLa *CCNF* K/O cells were cultured in DMEM (Sigma) containing 10% FBS (Gibco) and 100 U/ml penicillin, 100 mg/ml streptomycin (Millipore). HEK293, U-2-OS and HeLa cell lines were purchased from ATCC. *CCNF* K/O cells were generated previously from HeLa cells (Mavrommati et al, 2018). MCF7 were obtained from Ross Chapman laboratory. Cells were synchronized by addition of nocodazole 100ng/ml for 16h, mitotic cells were harvested by mitotic shake off and released to fresh media to allow progression into G1 phase where indicated.

### Antibodies and chemicals

Antibodies: CCNF (Santa Cruz), pRPA2 S4/8 (Bethyl), pRPA2 S33 (Bethyl), RPA2 (abcam), pKAP1 S824 (Genetex), KAP1 (Genetex), γ-H2AX (Millipore and Novus), pCHK1 S317 (Cell Signaling), CHK1 (), TFIIH (Santa Cruz), E2F1 (Santa Cruz), PCNA (Santa Cruz), histone 3, CCNE, CCNA (a kind gift of Michele Pagano), Exo1 (Genetex), SKP1 (Cell Signaling), pRRM2 T33 (D’Angiolella), RRM2 (Santa Cruz), pCDK substrate (Cell Signaling), GFP (Millipore), pHistone 3 S10 (Millipore), pp53 S15 (Cell Signaling), p53 (Santa Cruz), cdc6 (Santa Cruz), MTBP, Geminin (Santa Cruz), E2F3 (Santa Cruz), RRM1 (Santa Cruz), RRM2B (Santa Cruz), HA-HRP (Millipore).

Chemicals: ATRi (VE-821, Selleckchem), CHK1i inhibitors – UCN-01 (Selleckchem), LY2603618 (Selleckchem), CDK1 (Ro3306, Selleckchem), CDK2 (NU6140, Medchemexpress), EdU (Invitrogen), Calyculin A (Santa Cruz), cyclohexamide, MLN4924, L-PP (Santa cruz).

### Sample preparation: trypsin digestion

Cyclin F was eluted from the beads using 10mMTris, 2%SDS. The Cyclin F eluted fractions were sequentially incubated with DTT (5mM final concentration) and Iodacetamide (20mM final concentration) for 30min each at room temperature in the dark, before the proteins were precipitated twice with methanol/chloroform to remove SDS. Protein precipitates were reconstituted and denatured with 8M urea in 20mM HEPES (pH 8). Samples were then further diluted to a final urea concentration of 1M using 20Mm HEPES (pH8.0) before adding immobilised trypsin for 16h at 37°C (Pierce 20230). Trypsin digestion was stopped by adding TFA (final concentration of 1%) and trypsin removed by centrifugation. Tryptic peptides mixture was desalted using the SOLA HRP SPE cartridges (ThermoFischer) and dry down.

### LC_MS/MS and data analysis

Dried tryptic peptides were re-constituted in fifteen μl of LC-MS grade water containing 1% acetonitrile and 0.1% TFA. Thirty three percent of the sample was analysed by liquid chromatography-tandem mass spectrometry (LC_MS/MS) using a Dionex Ultimate 3000 UPLC coupled to a Q-Exactive HF mass spectrometer (Thermo Fisher Scientific). Peptides were loaded onto a trap column (PepMapC18; 300μm x 5mm, 5μm particle size, Thermo Fischer) for 1 min at a flow rate of 20 μl/min before being chromatographic separated on a 50cm-long EasySpray column (ES803, Thermo Fischer) with a gradient of 2 to 35% acetonitrile in 0.1% formic acid and 5% DMSO with a 250 nL/min flow rate for 60 min. The Q-Exactive HF was operated in a data-dependent acquisition (DDA) mode to automatically switch between full MS-scan and MS/MS acquisition. Survey-full MS scans were acquired in the Orbitrap mass analyser over an m/z window of 375 – 1500 and at a resolution of 60k (AGC target at 3e6 ions). Prior to MSMS acquisition, the top twelve most intense precursor ions (charge state >=2) were sequentially isolated in the Quad (m/z 1.2 window) and fragmented on the HCD cell (normalised collision energy of 28). MS/MS data was obtained in the orbitrap at a resolution of 30000 with a maximum acquisition time of 45ms, an AGC target of 5e5 and a dynamic exclusion of 27 seconds.

The raw data were searched against the Human UniProt-SwissProt database (June 2018; containing 20,361 human sequences) using Mascot data search engine. The search was carried out by enabling the Decoy function, whilst selecting trypsin as enzyme (allowing 1 missed cleavage), peptide charge of +2, +3, +4 ions, peptide tolerance of 10 ppm and MS/MS of 0.05 Da; #13C at 1; Carboamidomethyl (C) as fixed modification; and Deamidated (NQ), Oxidation (M), Phospho (ST), Phospho (Y) as a variable modification. MASCOT outputs were filtered using an ion score cut off of 20 and a false discovery rate (FDR) of 1%. A qualitative analysis was carried on with proteins identified with two or more peptides. The mass spectrometry raw data included in this will be deposited to the ProteomceXchange Consortium via the PRIDE partner repository.

### Kinase and drug screen

Hela and *CCNF* K/O cells were seeded at 1000 cells per well in 384-well plates 1 day before treatment with the KCGS library at 10uM, 1uM and 0.1uM concentrations. Relative cell survival was determined using resazurin 3 days after treatment. Z-scores were calculated for all samples and the difference between HeLa vs *CCNF* K/O cells was plotted.

### Relative cell proliferation assay

U-2-OS cells were seeded at 2000 cells per well to 96-well plate in hexaplicate 1 day before siRNA transfection. Cells were treated 1 day after siRNA transfection with indicated concentrations of ATRi or CHK1i. Hela and *CCNF* K/O cells were seeded at 2000 cells per well to 96-well plate in hexaplicate 1 day before treatment with indicated concentrations of ATRi or CHK1i. Relative cell proliferation was determined 3 days after treatment using resazurin 30ug/ml in complete media after 1-2h incubation.

### Cell counting and survival assay

U-2-OS cells were seeded at 20000 cells per well to 12-well plate 1 day before transfection with siRNA. Cells were treated 1 day after siRNA transfection with indicated concentrations of CHK1i and analyzed 3 days later. HeLa and *CCNF* K/O cells were seeded at 20000 cells per well to 12-well plate 1 day before treatment with treatment with 1uM CHK1i. Cells were counted and viability was determined using PI staining at 0.2ug/ml 24h, 48h and 72h by FACS Attune (Life Technologies) and data were analyzed using FlowJo software (TreeStar).

### Flow cytometry of chromatin bound proteins, cell cycle analysis and EdU staining

To analyze cell cycle profile, cell were pulse labelled with 10uM EdU for 30 minutes, harvested by trypsinization and fixed in 70% ethanol. For analysis of chromatin bound proteins, cells were pulse labelled with 10uM EdU for 30 minutes where indicated, harvested by trypsinization, washed in PBS, pre-extracted in pre-extraction buffer (25mM HEPES pH 7.4, 50mM NaCl, 1mM EDTA, 3mM MgCl2, 0.3M sucrose, 0.5% Triton X-100) 5 minutes on ice, centrifuged and fixed with 4% PFA 15 minutes at Room Temperature (RT). Fixed cell samples were permeabilized with 0.5% Triton X-100 for 15 minutes and blocked in 1% BSA. Click reaction was performed in buffer containing 0.1M Tris-HCl pH 8.5, 0.1 M sodium ascorbate, 2mM CuSO4, 10uM Alexa647-azide 30 minutes at room temperature. Cells were washed in PBS and incubated with primary antibodies for 2h at RT before incubation with secondary antibodies 1h at RT. Cells were resuspended in DAPI 5ug/ml in PBS and data were acquired using FACS Attune (Life Technologies) and analyzed using FlowJo (TreeStar).

### Immunoprecipitation and immunoblotting

HEK293 cells were seeded to 10cm plates 1 day before plasmid transfection (5ug/plate using PEI-MAX 40kDA at ratio 6:1). After 20h, cells were treated with 1uM MLN4924 or 1uM CHK1i for 4-5h. Cells were washed twice with PBS, harvested to lysis buffer (50mM Tris pH 7.5, 150mM NaCl, 10mM β-glycerophosphate, 10% glycerol, 1% Tween-20, 0.1% NP-40) containing protease inhibitors (Sigma), beta-glycerolphosphate and okadaic acid, incubated on ice for 30minutes and centrifuged for 30 minutes at 20 000g 4C. Supernatant was incubated at 4C with either GPF-Trap beads or HA-beads for 1h and 2h, respectively, before washing 4 times with lysis buffer. To dephosphorylate IPs, beads were incubated with λ-phosphatase in appropriate buffer for 30 minutes at 30°C with orbital shaking and washed 3 times with lysis buffer. Immunoprecipitates were eluted using 2x LDS buffer (Life Technologies) supplemented with β-mercaptoethanol and boiled for 5min at 95°C. Whole cell lysates were harvested using 2xSB (625mM Bis-Tris pH 6.8, 20% (w/v) glycerol, 4% (w/v) SDS), boiled, sonicated and protein concentration was evaluated using BCA protein kit (Thermo) in most cases. Cell lysate or immunoprecipitate was resolved in 10% Bis-Tris gels, transferred to nitrocellulose membrane (Millipore) and immunoblotted.

### DNA constructs, siRNA and transfections

CCNF gene was subcloned from Flag-Cyclin F(D’Angiolella et al, 2012) into pYFP-C1. The product YFP-cyclin F was fully sequenced verified. HA-E2F1 plasmid was a kind gift of Nick La Thangue. Plasmid transfections were carried using PEI (PEI-MAX 40kDA, Polysciences) at ratio 6:1.

All siRNAs were purchased from Ambion: NC (4390843), CCNF (s2527, s2528, s2529), cdc6 (s2744), Cdt1 (s37724), Geminin (s27307), MTBP (s25788), TICRR (s40361), E2F3 (s4411), E2F1 (s4405, s4406), Exo1 (s17502) and transfected 10nM. siRNA transfections were performed using HiPerfect (Qiagen) following manufacturer instructions.

### Comet assay

Alkaline and neutral comet assays were performed in parallel on the same cell set. Per condition, 50,000 trypsinised cells were mixed with 1% low melting point agarose (37 °C) and added to microscope slides pre-coated with 1% normal melting point agarose. For the alkaline assay: cells were lysed for 1 h at 4°C in a buffer containing 2.5 M NaCl, 100 mM EDTA, 10 mM Tris-HCl, 1% (v/v) DMSO and 1% (v/v) Triton X-100, pH 10.5. Slides were placed in an electrophoresis tank and covered with 4°C electrophoresis buffer (300 mM NaOH, 1 mM EDTA, 1% (v/v) DMSO, pH>13) for 30 m, to allow DNA to unwind. Electrophoresis was performed at 25 V for 25 m at 300 mA. Slides were neutralised in 0.5 M Tris-HCl pH 8.1 and allowed to dry overnight. For the neutral assay: cells were lysed for 2 h at 4°C in a buffer containing 2.5 M NaCl, 100 mM EDTA, 10 mM Tris-HCl, 1% (v/v) sodium lauryl sarcosinate, 10% (v/v) DMSO and 0.5 % (v/v) Triton X-100, pH 9.5. Slides were washed in TBE buffer (4°C) and electrophoresis was performed in TBE buffer (4°C) for 25 m at 25 V. Slides were allowed to dry overnight. For both assays, cells were rehydrated in H2O, stained with SYBR Gold for 30 m, and allowed to dry overnight. Analysis was performed using the Nikon NiE microscope and Andor Komet7.1 software. Two microscope slides were analysed per condition, with at least 100 cells per condition per biological replicate. Data were expressed as Olive tail moment: (Tail.mean - Head.mean)*Tail%DNA/100. p-values were calculated using the two-tailed Mann-Whitney test.

### Gene set enrichment analysis

Pathway Analysis was done using Benjamini-Hochberg corrected Hypergeometric test as applied in GeneCodis3(Tabas-Madrid et al, 2012). Gene Ontology Biological Processes (BPs) were considered. The top 50 gene were included in the analysis and BPs with an adjusted significance level of 0.002 or lower were considered significant.

### Statistical analysis

To evaluate statistical significance p-value was determined using GraphPad Prism. Paired two-tailed t-test was used for relative survival assays. Two-way anova test was used to evaluate statistical significance of cell proliferation in cell counting assay. Unpaired two-tailed t-test was used to evaluate statistical significance of percentage of γ-H2AX and hyperRPA positive cells. p-values are marked by asterisks in figures as follows: * p<0.05, ** p<0.005, *** p<0.0005.

## Expanded View Figure Legends and Table 1

**Expanded View Figure 1. Related to Figure 1.**
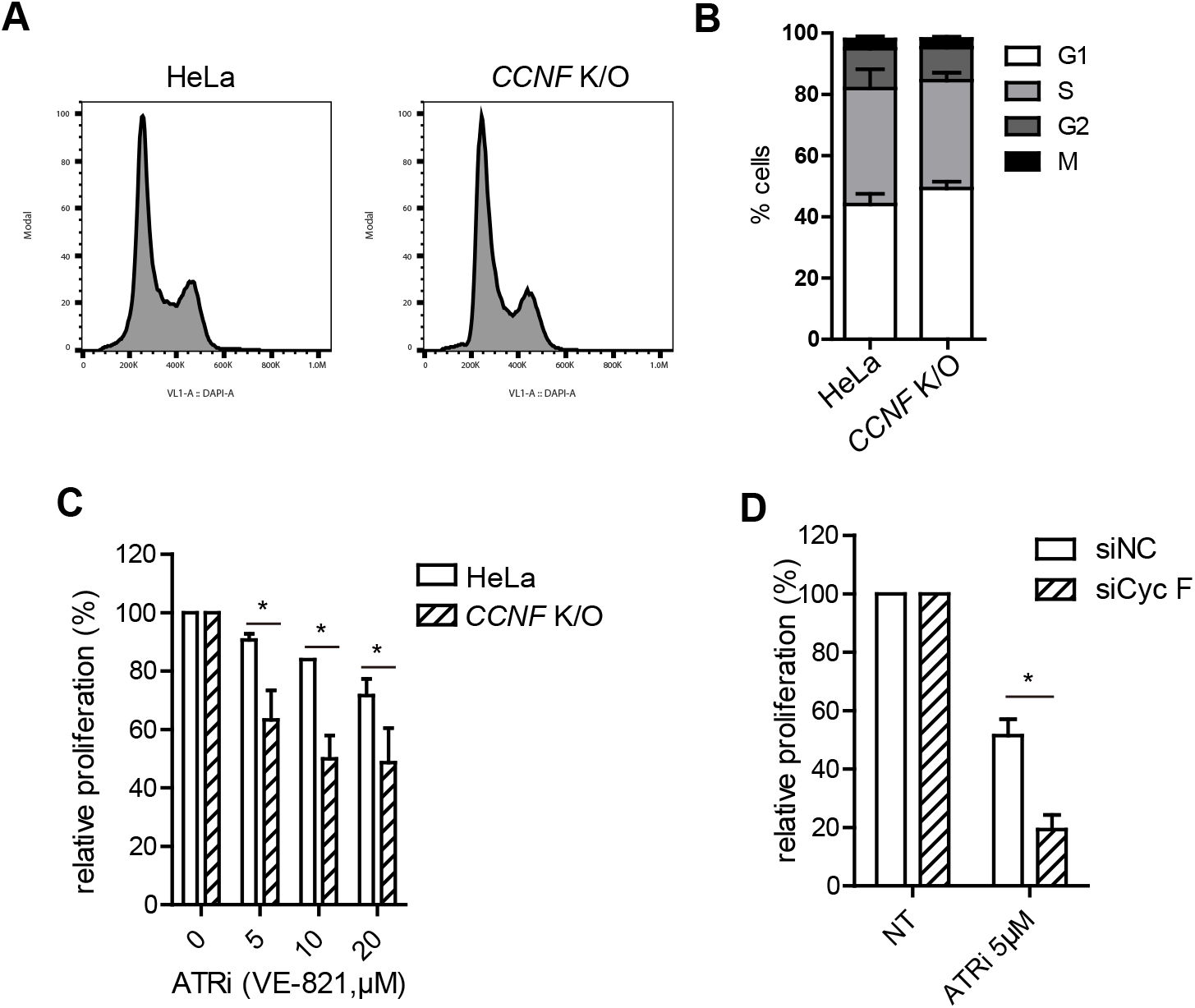
A. Cell cycle profile of HeLa and CCNF K/O detected by FACS using Propidium Iodine (PI). B. Cell cycle distribution of untreated HeLa and *CCNF* K/O cells using combined phospho Histone H3 Serine 10 and EdU staining. C. Cell survival measured using rezasurin and compared to controls treated with DMSO (expressed as relative proliferation %). Cells were treated with ATR inhibitors at the indicated concentrations. Data were plotted with ± SD (*<0.05 p-value). D. Cell survival of U-2-OS cells transfected with non targeting siRNA siNC (Negative Control) or siCyc F after treatment with Chk1i (LY2603618) at indicated concentrations compared to DMSO treated controls (NT). Data were plotted with ± SD (*<0.05 p-value).

**Expanded View Figure 2. Related to Figure 2.**
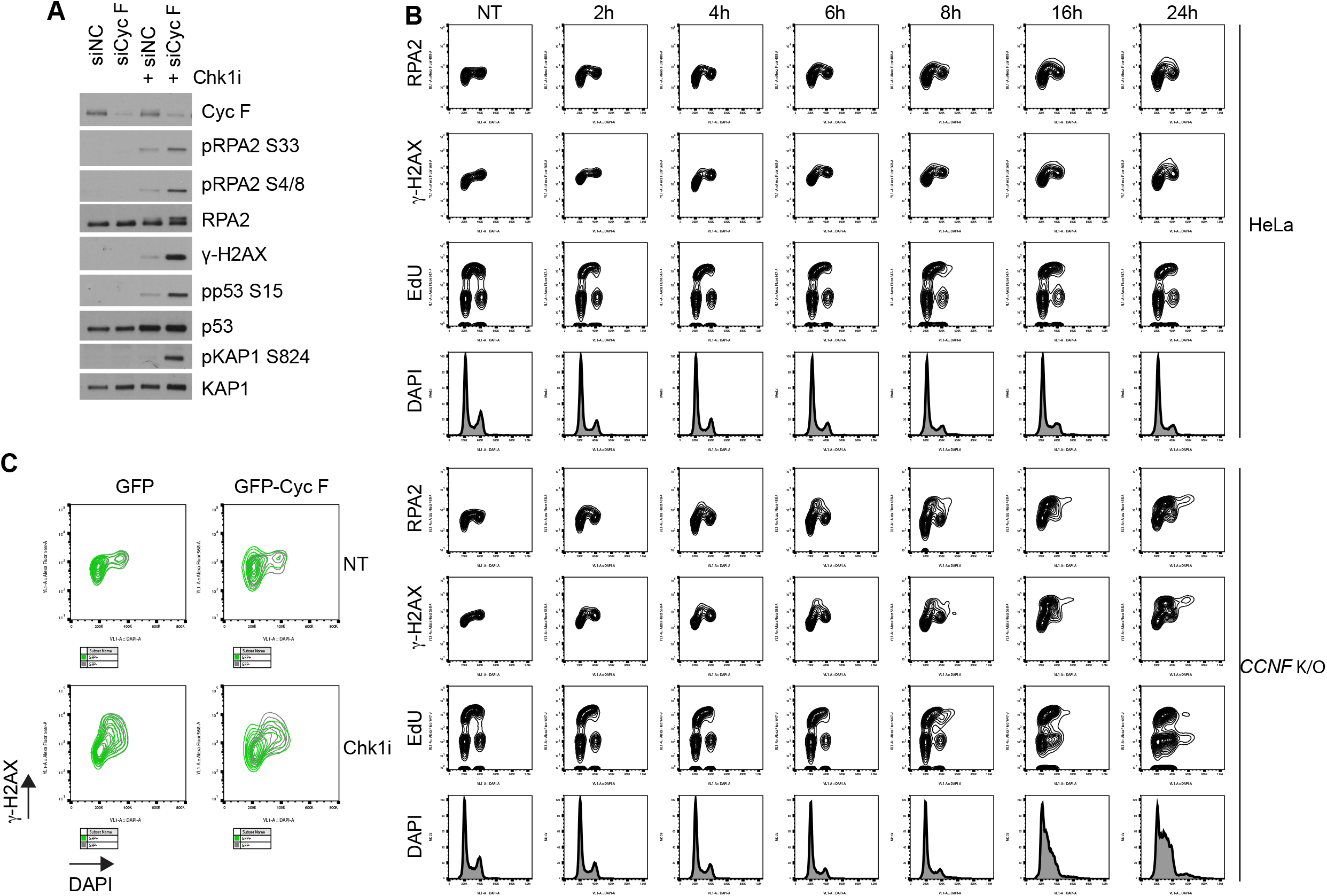
A. U-2-OS cells transfected with the indicated siRNA and treated with Chk1i (LY2603618) for 20 hours were harvested and lysed using SDS. Indicated protein were resolved by SDS page and detected by Western blot (Wb). B. Representative plots of HeLa and *CCNF* K/O cells treated with 1μM Chk1i for the indicated hours (h). Half an hour before harvesting, cells were incorporated with EdU (10uM as final concentration). EdU by click reaction, γ-H2Ax and RPA by immunostaining, and DAPI staining were detected by FACS analysis. C. Representative plots of γ-H2AX intensity vs DAPI signal of *CCNF* K/O cells transfected with empty GFP and YFP-CCNF untreated (*top panels)* and after treatment with 1μM Chk1i (LY2603618) for 20h (*bottom panels*).

**Expanded View Figure 3. Related to Figure 3.**
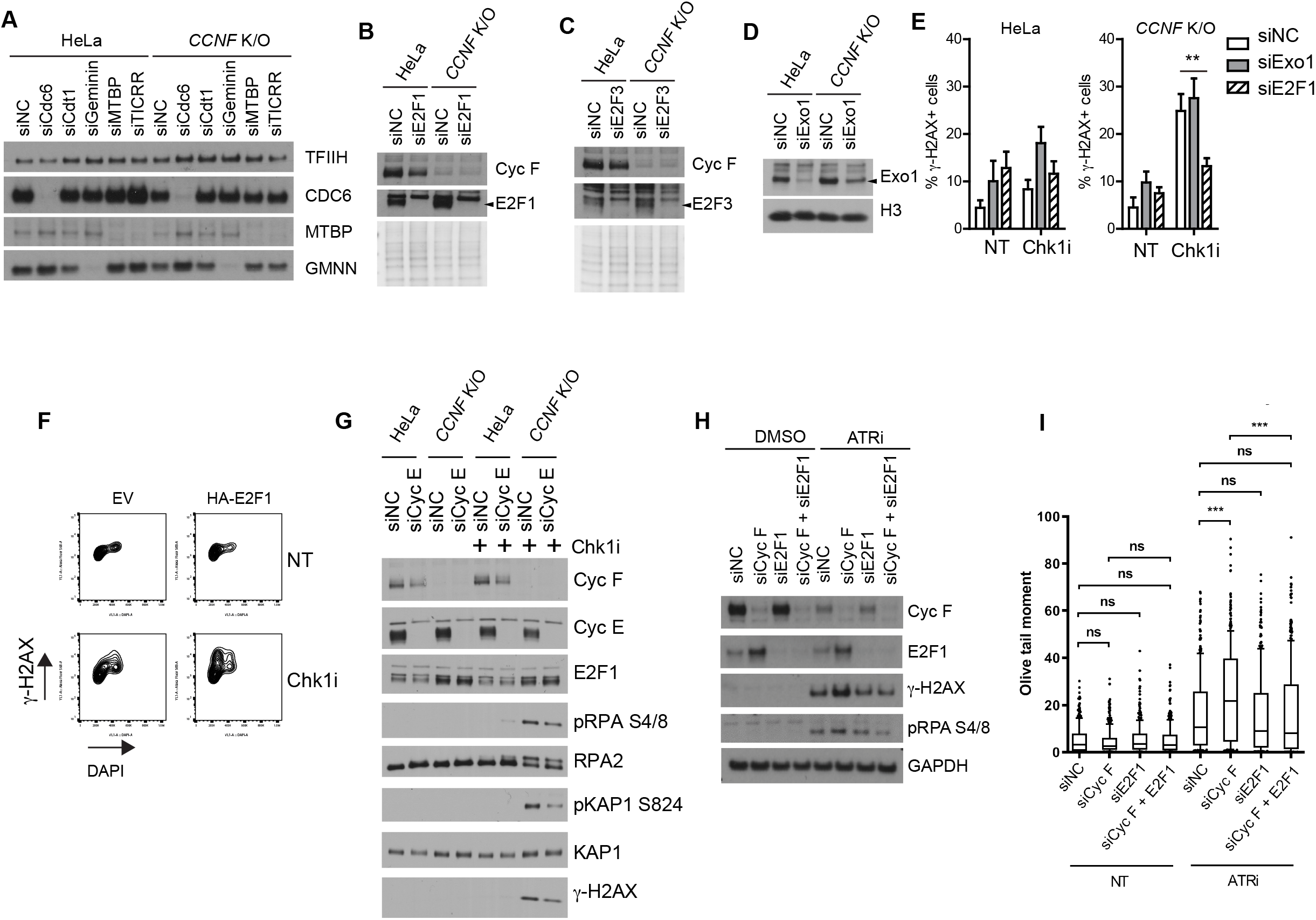
A. HeLa and *CCNF* K/O cells transfected with the indicated siRNA for 48 hours were harvested and lysed using SDS. Indicated protein were resolved by SDS page and detected by Wb. The image verifies knockdown efficiency of experiments presented in Figure 3B and C. TFIIH was used as a loading control. B. Wb of indicated proteins as in A. Ponceau S staining is a loading control. C. Wb of indicated proteins as in A. Ponceau S staining is a loading control. D. Wb of indicated proteins as in A. H3 served as a loading control. E. Percentage (%) of γ-H2AX positive cells in HeLa and *CCNF* K/O after transfection of the indicated siRNAs treated with Chk1i for 20 hours. Percent of γ-H2AX positive cells were plotted as Mean % ± SD (**< 0.005 p-value). F. Representative FACS plots of γ-H2AX *vs* DAPI signal of HeLa cells transfected with Empty Vector (EV) and HA-E2F1 left untreated or after treatment with 1μM Chk1i for 20h. G. HeLa and *CCNF* K/O cells transfected with the indicated siRNA and treated with 1μM Chk1i (LY2603618) for 20 hours were harvested and lysed using SDS. Indicated protein were resolved by SDS page and detected by Wb. H. Cells transfected with non targeting siRNA (siNC) and indicated siRNA were left untreated (DMSO) or treated with ATRi (VE-821; 1μM) for 24 hours. Cells were harvested and lysed using a triton based extraction buffer. Indicated protein were resolved by SDS page and detected by Wb. I. Quantification of comets in alkaline gels. At least 100 cells, across two slides, were analysed in each condition in three biological replicates. Data are shown as medians, with 25/75% percentile range (box) and 10-90% percentile range (whiskers). Fold changes of median values are shown in bold. p-values were calculated using the Mann-Whitney test (two-tailed). Olive tail moment = (Tail.mean - Head.mean)*Tail%DNA/100.

**Expanded View Figure 4. Related to Figure 4.**
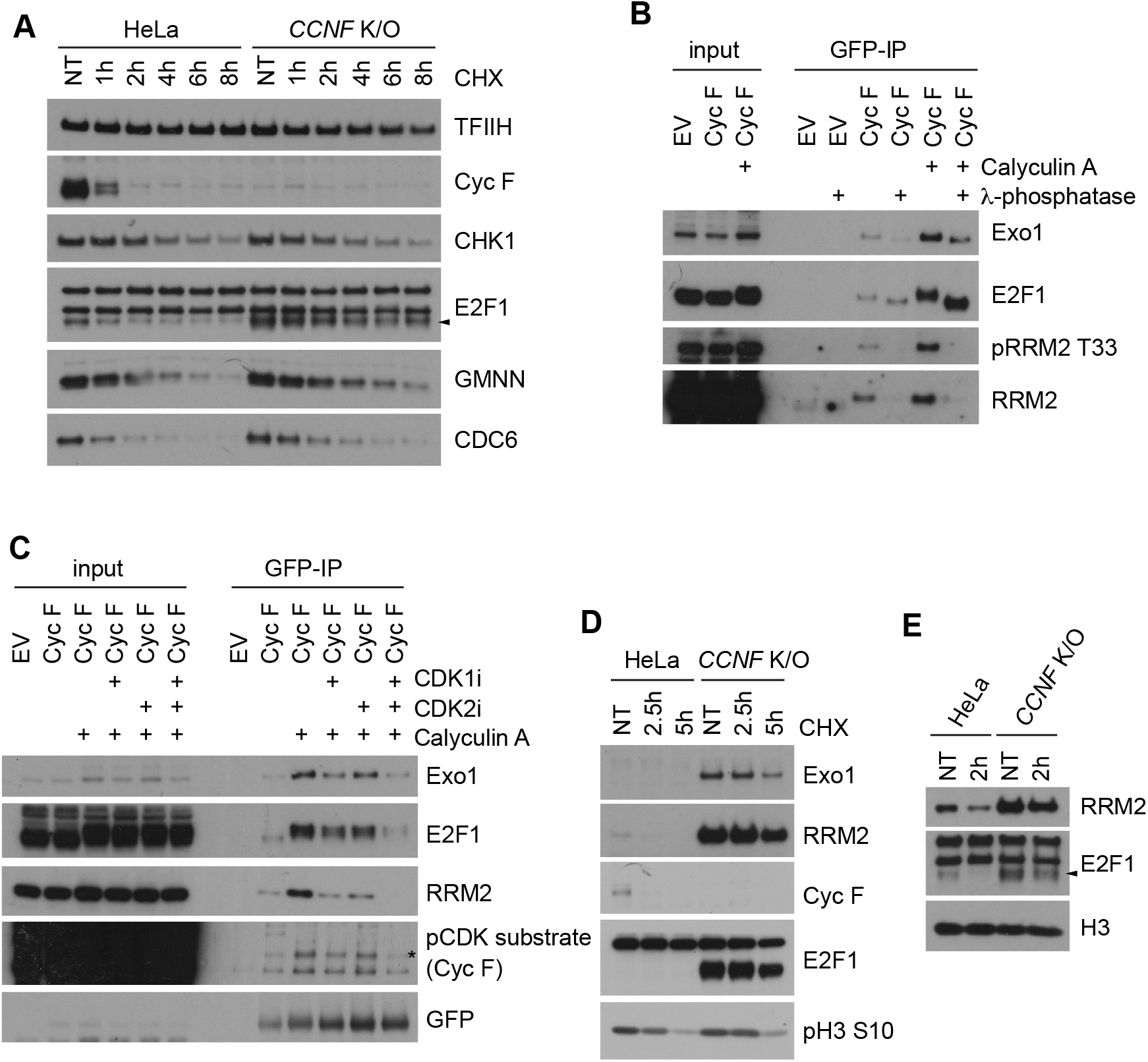
A. HeLa and *CCNF* K/O cells were treated with CycloHeximide (CHX-50ug/ml) for the indicated hours (h), harvested and lysed using SDS. Indicated protein were resolved by SDS page and detected by Wb. TFIIH was used as a loading control. B. HEK293 cells transfected with Empty Vector (EV-GFP) or YFP-Cyclin F were treated with Calyculin A(15 minutes-+) where indicated. Harvested cells were processed for immunoprecipitation using GFP-nanobodies. Immunoprecipitates were treated with λ-phosphatase as indicated (+), and immunoblotted as indicated. C. HEK293 cells transfected with Empty Vector (EV-GFP) or YFP-Cyclin F were treated with Calyculin A (15 minutes-+), CDK1 and CDK2 inhibitors where indicated(+). Harvested cells were processed for immunoprecipitation using GFP-nanobodies. Immunoprecipitates were immunoblotted as indicated. D. HeLa and *CCNF* K/O cells synchronized in mitosis by nocodazole (100ng/ml for 16h) were treated with CycloHeximide (CHX-50ug/ml) for the indicated hours (h). Indicated protein were resolved by SDS page and detected by Wb. E. HeLa and *CCNF* K/O cells synchronized in mitosis by nocodazole (100ng/ml for 16h) were released from Nocodazole block in fresh media for the indicated hours (h). Indicated protein were resolved by SDS page and detected by Wb.

